# Elucidating the half-site reactivity mechanism of *Salmonella enterica* FraB deglycase using native mass spectrometry

**DOI:** 10.64898/2026.07.17.739170

**Authors:** Yuan Gao, Jamison D. Law, Venkat Gopalan, Vicki H. Wysocki

## Abstract

Inter-subunit communication and allosteric regulation are central to the function of oligomeric enzymes, yet these features remain difficult to characterize. Conventional kinetic and structural methods typically yield ensemble averages or static snapshots, thus making it difficult to uncover the dynamic cross-subunit cooperation obligatory for multi-site catalysis by oligomeric enzymes. Here, we investigate *Salmonella* FraB—a homodimeric deglycase and a potential drug target—to showcase the value of an integrated approach combining native mass spectrometry (nMS), surface-induced dissociation (SID), and kinetic studies to gain insights into catalytic intermediates and inter-subunit communication. By resolving substrate-, product-, and mixed-occupancy species, nMS revealed that both inter-subunit active sites in FraB bind substrate even though only one catalytic center generates the product at any given time. To characterize each active site independently, we designed heterodimers with a mutation that changes the general base or acid in only one active site. Kinetic studies with these mutants indicate that although the two active sites are likely coupled, they do not concomitantly perform cleavage. Consistent with the conformational asymmetry observed in *apo*-FraB crystal structures, our findings establish a half-site reactivity mechanism in which post-binding conformational changes across the dimer interface restrict substrate cleavage to one active site even though both protomers are able to bind substrate. Importantly, this nMS-based workflow offers a broadly applicable framework for resolving the catalytic states and inter-site communication of oligomeric enzymes that are otherwise difficult to uncover by conventional structural methods.

## Introduction

The majority of protein enzymes function as oligomers, with nearly one-half adopting a dimeric structure.^1^ Understanding the coordination between the multiple active sites in these oligomers is necessary to establish their catalytic mechanisms, deduce the evolutionary driving forces for oligomeric enzymes, and formulate principles to design superior biocatalysts. Traditional enzyme assays provide bulk turnover rates, but do not reveal catalytic performance at the individual active sites. High-resolution structural biology methods have had variable success in uncovering the structural reorganization during catalysis. Standard X-ray crystallography and cryo-EM approaches often provide static snapshots and therefore may not directly reveal real-time conformational changes.^2,3^ Time-resolved serial and freeze-trapping crystallography as well as “mix-and-spray” cryo-EM are beginning to reveal exquisite details of the distinct molecular species along the catalytic cycle. However, their broader application can be limited by the need for protein microcrystals, precise reaction synchronization, and complex data interpretation.^2,4,5^ Nuclear magnetic resonance (NMR) spectroscopy is a powerful tool for studying protein dynamics, but it is limited by protein size and requires a large amount of sample.^6,7^ Here, we demonstrate the value of native mass spectrometry (nMS) to dissect functional cooperation in dimeric enzymes.

nMS has emerged as a powerful structural biology method for studying enzyme complexes because it retains non-covalent interactions and preserves native-like structures of the analytes.^8–10^ nMS enables direct observation of protein-ligand complexes,^11,12^ including enzyme-substrate-inhibitor complexes.^13–15^ nMS can resolve individual ligand-bound enzyme species by mass, allowing the species of interest to be isolated in the gas phase and further characterized using activation techniques. Collision-induced dissociation (CID) is the most commonly used dissociation method, in which protein ions collide multiple times with gas molecules and undergo structural rearrangements prior to complex dissociation and ligand loss. In comparison, surface-induced dissociation (SID), a method in which ions undergo a single, rapid collision with a solid surface, minimizes structural rearrangement prior to dissociation, making it a useful tool for maintaining native-like subunit structure and localizing ligand binding.^16–19^

Here, we used nMS to study *Salmonella enterica* FraB, a homodimeric deglycase. In the final step during the catabolism of fructose-asparagine, FraB cleaves 6-phosphofructose-aspartate (6-P-F-Asp, 375 Da) to glucose-6-phosphate (G-6-P, 260 Da) and aspartic acid (Asp, 133 Da) (Figure 1A). Because the deletion of *fraB* causes a build-up of 6-P-F-Asp and thereby intoxicates *Salmonella*, FraB is a promising drug target.^15,20,21^ Homology modeling of the FraB homodimer and the docking of 6-P-F-Asp revealed two inter-subunit active sites at the dimer interface, each consisting of catalytic residues from opposing protomers. Previous studies demonstrated that E214 and H230 serve as the general base and acid, respectively, during the deglycation reaction (Figure 1A, Figure S1).^21^ The E214A and H230A FraB mutants are inactive *in vitro*; moreover, introducing these mutations individually in the *fraB* gene of the *Salmonella* chromosome abolished the pathogen’s ability to grow on fructose-asparagine. The recent high-resolution *apo*-FraB crystal structure revealed an unexpected conformational asymmetry, with the two chemically identical subunits adopting distinct conformations.^22^ Because co-crystallization with the substrate has been elusive, it remains unclear if the “conformational heterodimer” results in two disparate active sites. Here, by combining nMS, SID and kinetic analysis, we gained insights into the mechanism and the structural changes that occur during catalysis by the FraB homodimer. This strategy should be broadly applicable to other dimeric and higher-order enzyme oligomers whose mechanisms involve conformational restructuring and communication between active sites.

**Figure 1.**
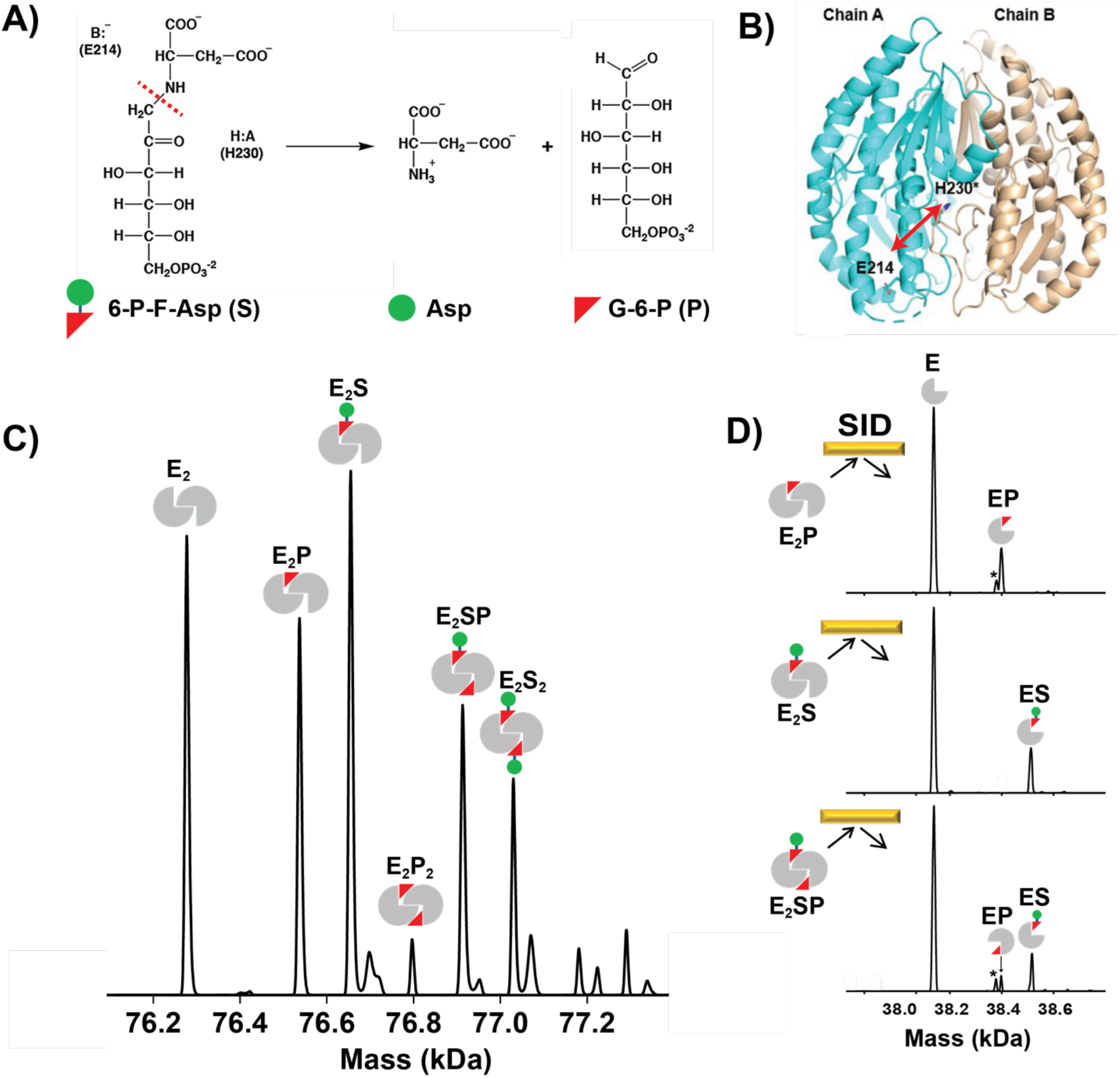
(A) Proposed catalytic mechanism for FraB using H230 and E214 as a general acid and base, respectively (adapted from Sengupta et al., 2019).^21^ (B) Crystal structure of *apo*-FraB dimer showing the separation of E214 and H230 across the subunit interface (adapted from Zakharova et al., 2025).^22^ (C) Deconvolved native MS spectrum of FraB in the presence of 6-P-F-Asp substrate. (D) Isolation followed by SID of E_2_P, E_2_S, and E_2_SP. The asterisk (*) represents peaks corresponding to loss of a H_2_O molecule from G-6-P.

## Results

### nMS with the FraB homodimer: Implications for catalysis

The conformational asymmetry in the FraB dimer might reflect non-equivalent active sites that undergo correlated rearrangements as part of the molecular choreography central to catalysis: one site might close to bind the substrate, while opening of the second site releases the product. nMS is particularly well suited to test this temporal sequence of events with FraB, especially given its successful use in showing that the FraB E214A and H230A mutants can bind 6-P-F-Asp but not catalyze substrate cleavage or product formation.^21^

The nMS data for the FraB homodimer with its substrate, 6-P-F-Asp (2 µM FraB:300 µM 6-P-F-Asp) resolve a series of enzyme species differing in ligand stoichiometry (Figure 1C; nMS spectrum for FraB alone shown in Figure S2). Substrate-bound species E_2_S and E_2_S_2_ are observed, consistent with 6-P-F-Asp bound to the two active sites. Moreover, we detect E_2_SP, which corresponded to one substrate and one product (G-6-P) bound to the FraB homodimer simultaneously, suggesting that substrate binding and product formation can occur concomitantly and independently at the two active sites. These findings support three inferences. *First*, the FraB homodimer is catalytically active prior to electrospray ionization, and both the substrate- and product-bound enzyme complexes are kinetically trapped during the short timescale of the nMS experiment, with their non-covalent interactions preserved. *Second*, we can rule out product release as mandatory for substrate binding by the adjacent subunit. *Last*, despite formation of two products (G-6-P and Asp), we detected FraB bound only to G-6-P, a finding that we attribute to differences in affinity of FraB for the two products. The selective binding of G-6-P over Asp suggests that FraB binds to sugar rather than the amino acid in 6-P-F-Asp. This notion is consistent with the homology model, in which docking of 6-P-F-Asp predicts that the phosphate interacts with residues S32, E85, and Y325 in FraB.^21^ Direct incubation of FraB with free G-6-P produced multiple product-bound species, indicating that G-6-P can bind nonspecifically to FraB. Additional inhibitor-blocking experiments further support this assignment (see Figure S3 for additional details). Thus, the minor E_2_P_2_ population observed by native MS (∼4%; Figure 1C) is most consistent with nonspecific product rebinding rather than simultaneous dual-site catalysis.

Each ligand-bound species, E_2_S, E_2_P, and E_2_SP, was isolated by the quadrupole from the heterogeneous mixture and characterized by gas-phase activation. CID dissociated the substrate or product before the dimer interface was disrupted, providing little subunit-level information. In comparison, SID cleaves the dimer along the monomer–monomer interface and partially retains ligands bound at that interface (Figure 1D). E_2_S dissociated into E and ES; E_2_P dissociated into E and EP. For E_2_SP, we observed all three predicted monomer species (E, ES, and EP), suggesting that one subunit was bound to the substrate while the neighboring protomer bound the product. Although some ligand-bound monomers dissociated to *apo*-monomer due to the activation, the retention of ligand on the released monomers demonstrates the utility of SID for identifying the ligand bound to each subunit. Collectively, these data support the idea that FraB functions as a dimer with two distinct active sites capable of independently binding substrate and forming product.

### Kinetic studies with FraB heterodimers

The crystal structure and homology model establish that each FraB active site is assembled from E214 of one subunit and H230 of the other, forming two interlocking catalytic pockets at the dimer interface.^21,22^ As any point mutation introduced into the FraB monomer will be present in both catalytic centers of the homodimer, we designed FraB heterodimers in which an active site mutation is restricted to one site, leaving one wild-type and one mutated catalytic pocket within the same dimer. FraB heterodimers were generated by co-overexpressing two different affinity tagged FraB variants: His_6_-FraB and PA14-FraB. Our initial plan was to leverage the His_6_ and PA14 tags for tandem immobilized metal- and dextran-binding^23^ affinity chromatography. However, sequential dextran-affinity and size-exclusion chromatography sufficed to isolate FraB heterodimers with distinct active sites (Figure S4).

We denote subunits as E (wild type), E_e_ (E214A), or E_h_ (H230A), and the four heterodimer variants as: PA14-FraB:His_6_-FraB (EE; completely wild-type); PA14-FraB:His_6_-FraB E214A (EE_e_); PA14-FraB:His_6_-FraB H230A (EE_h_); and PA14-FraB E214A:His_6_-FraB H230A (E_e_E_h_) (Figure 2A). SDS-PAGE analysis revealed the successful purification of PA14-FraB (58 kDa) and His_6_-FraB (38 kDa) (Figure S6). To verify heterodimer formation and accurate mass, native MS experiments were performed on each heterodimer variant. The dominant species observed were heterodimers as expected, with minor populations of homodimer (Figure S5). SID of isolated heterodimers generated the expected monomer products corresponding to the His_6_- and PA14-tagged subunit variants, confirming the heterodimer composition of the parental species. All measured masses matched the expected masses (< 2 Da error).

**Figure 2.**
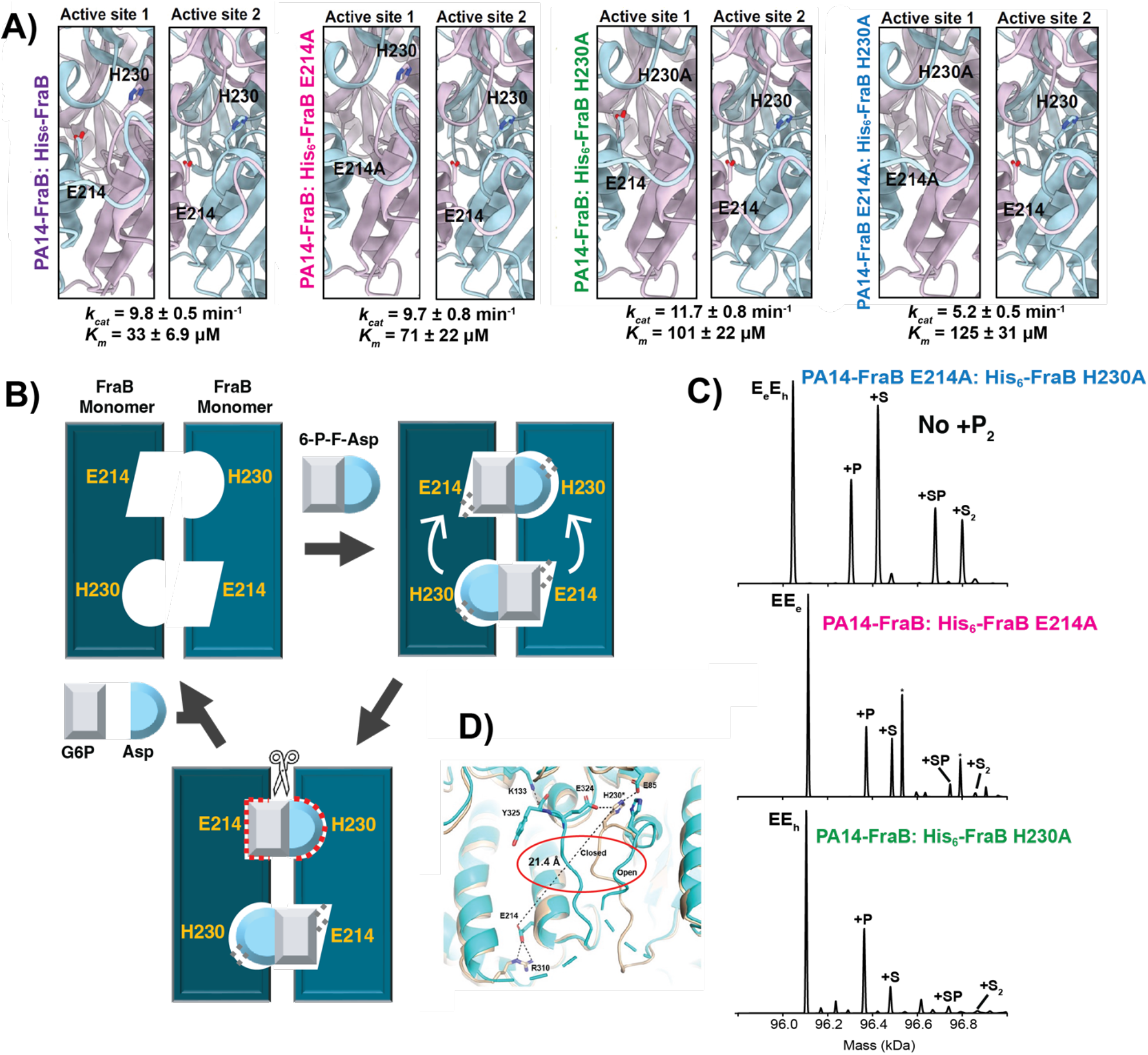
Heterodimer mutants support half-site reactivity in the FraB dimer. (A) Design of FraB heterodimer active-site mutants and corresponding kinetic parameters (see supplement for the primary kinetic data). (B) Schematic illustrating FraB half-site reactivity. (C) Deconvolved native MS spectrum of FraB heterodimer variants in the presence of 6-P-F-Asp, showing formation of E_2_S and E_2_S_2_ species while product-containing complexes remain limited to single-site cleavage, with no detectable E_2_P_2_. (D) Crystal structure of the *apo*-FraB homodimer with overlay of the two active sites that depicts their structural plasticity (adapted from Zakharova et al., 2025).^22^

Kinetic studies with the FraB heterodimer variants revealed unexpected catalytic behavior (Figure 2A; see Figure S6 for Michaelis–Menten data). Mutating either E214 or H230 to alanine at a single active site in the FraB homodimer (EE_e_ and EE_h_) did not significantly affect the *k*_*cat*_, although the *K*_*m*_ increased 2- to 3-fold. In contrast, when both E214A and H230A mutations were introduced (E_e_E_h_) to the same inter-subunit active site, the *k*_*cat*_ and *k*_*cat*_/*K*_*m*_ were reduced by 2- and 7-fold, respectively. Because E214A or H230A FraB homodimers are catalytically inactive, the retention of wild-type-like *k*_*cat*_ despite selective mutation of E214 or H230 in a single active site suggests that only one active site is typically functional during a single catalytic cycle. Moreover, E_e_E_h_ has a 2-fold lower *k*_*cat*_ compared to EE_e_ and EE_h,_ although all three mutant variants have one inactive site (Figure 2A), suggesting that the two sites are coupled and do not act independently. These findings support the idea of half-site reactivity during FraB catalysis.

### nMS studies with FraB heterodimers

To further examine how site-specific mutation in individual active sites affects ligand occupancy, we analyzed the heterodimer variants by nMS in the presence of 6-P-F-Asp (Figure 2C). All three variants (EE_e_, EE_h_, and E_e_E_h_) have E_2_S_2_ and E_2_SP, indicating that both active-site pockets remain capable of substrate binding despite mutation of key catalytic residue(s). Product-containing species were also observed, primarily as E_2_P and E_2_SP, whereas E_2_P_2_ was not detected. While the absence of E_2_P_2_ is expected for heterodimers containing an inactive active site, the coexistence of E_2_S_2_ and E_2_SP indicates substrate can occupy both sites, but product formation is restricted to one site, and that substrate- and product-bound species can coexist within the same dimer. [An additional peak in the EE_e_ spectrum (asterisk, 419 Da) corresponds to a 6-P-F-Asp– ethylene glycol adduct (+44 Da, ethylene glycol with loss of water), a buffer-derived contaminant.]

To further characterize ligand localization within the heterodimer, the E_e_E_h_ complex was isolated and subjected to SID (Figure S7). Conventional CID of the heterodimer resulted primarily in ligand dissociation without monomer dissociation. In contrast, SID of the isolated heterodimer complex generated dissociated PA14-FraB and His_6_-FraB monomers with partially retained substrate and/or product binding. Because FraB catalysis occurs at inter-subunit active sites, ligand retention on both subunits is consistent with binding at the dimer interface. Notably, PA14-FraB E214A, which has a functional H230, showed more ligand retention, including substrate- and product-bound species and a minor ESP complex (Figure S7). In comparison, the His_6_-FraB H230A subunit exhibited reduced ligand retention, highlighting the greater importance of H230. These observations are consistent with the X-ray crystal structures showing large-scale movements of the H230-containing loop as part of the switch from open to closed conformations, which in turn dictates ligand binding.^22^ Overall, the SID results provide subunit-level evidence that ligand interactions are distributed across the dimer interface and that both subunits contribute to ligand binding and catalysis.

## Discussion

Examples of half-site reactivity were established from studies that investigated kinetics, binding stoichiometry, or negative cooperativity in oligomeric enzymes.^1,24–29^ This mechanistic theme highlights how ligand binding at one site in an oligomeric enzyme restructures the energy/conformational landscape to decrease ligand interactions and function at a second site. Our use of nMS to obtain evidence for substrate-, product-, and mixed-occupancy bound species during catalysis by the FraB deglycase homodimer establishes the utility of this method in mechanistic enzymology. A collective assessment of our findings provides additional insights into FraB catalysis.

The near-absence of E_2_P_2_ in our nMS data confirms the non-equivalence of the two active sites in FraB, as anticipated from the high-resolution structures. However, as with alternating pistons in an engine, it is possible that one site in FraB traps the substrate, even while the simultaneous opening of the second site releases the product. Nevertheless, the simultaneous detection of E_2_S, E_2_S_2_, E_2_P, and E_2_SP species by nMS demonstrates that both FraB active sites can concurrently bind the substrate. Furthermore, the mixed-occupancy E_2_SP intermediate reveals that substrate and product can coexist in the same dimer. If there is concomitant occupancy of substrate at both sites, one could envision active-site switching after each cleavage but this idea is not supported by our kinetic data. With E214A or H230A at a single active site in the FraB homodimer (EE_e_ or EE_h_), the *k*_*cat*_ was unaffected (Figure 2, Figure S6). Nevertheless, an alternating catalysis model cannot be entirely ruled out if chemistry is rate-limiting and substrate association kinetics are rapid; in this case, an incapacitated second site is unlikely to lead to a lower turnover rate.

A more likely scenario for FraB catalysis might mirror the widespread behavior documented in homodimeric RNA modifying enzymes (e.g., adenosine deaminase),^30,31^ where one subunit is involved in substrate recognition and the other performs catalysis. Such a model is consistent with our finding of a 2-fold decrease in *k*_*cat*_ and a 7-fold decrease in *k*_*cat*_ /*K*_*m*_ when E214A and H230A mutations are present in one active site (E_e_E_h_). Thus, substrate binding and cleavage in one site are clearly perturbed by changes to the second site. nMS data show that the E_e_E_h_ mutant can bind substrates at both sites, but the microenvironment at the mutated site is significantly altered, failing to trigger the conformational restructuring necessary for optimal catalysis by the second site, which has the general acid and general base. Taken together, our data indicate that post-binding of 6-P-F-Asp to both sites in the FraB homodimer, one site positions E214 and H230 for optimal cleavage largely due to a conformational rearrangement engendered by the second site (Figure 2B). Cleavage releases Asp but leaves G-6-P transiently bound to the second site, as evidenced by the mixed-occupancy E_2_SP species detected by nMS (Figure 1C).

While half-site reactivity might seem an evolutionary indulgence, this feature could provide payoffs. With FraB, tasking one site with substrate recognition prior to licensing cleavage by the second site is an allosteric regulation safeguard that minimizes promiscuous activity on other cellular sugar-phosphates. The FraB crystal structure revealed how the ∼21 Å distance between E214 and H230 in the ground state (incompatible with catalysis) is decreased to 7.5 Å by rearranging the E214-containing helix from a normal (i+4) to a Pi (i+5) configuration.^22^ We postulated that binding the correct substrate in one inter-subunit active site triggers the normal to Pi helix shift and promotes a cleavage-competent state in the second catalytic center. This conceptual framework should inform future studies aimed at delineating additional elements of the FraB half-site reactivity mechanism. More broadly, this work demonstrates the capability of native MS to resolve multi-site substrate and product distributions during enzyme catalysis and to provide mechanistic insights that are difficult to obtain by other methods.

## Author Contributions

Yuan Gao: Writing – review and editing; original draft; investigation; methodology. Jamison D. Law: Writing – review and editing; investigation; methodology. Venkat Gopalan: Investigation; review and editing; methodology; funding acquisition; supervision; conceptualization. Vicki H. Wysocki: Investigation; review and editing; methodology; funding acquisition; supervision; conceptualization.

## Acknowledgements

This work was funded by grants from the National Institutes of Health (P41GM128577 and RM1GM149374 to V.H.W.; R01AI140541 and R01AI116119 to V.G., and V.H.W.; T32-GM086252 to JDL, a Pelotonia Graduate Fellowship to J.D.L)

