## Supporting Information for "Elucidating the half-site reactivity mechanism of *Salmonella enterica* FraB deglycase using native mass spectrometry"

### Methods

#### FraB heterodimer overexpression and purification

To enable co-expression of pLANT-2b–$His_6$-TRS-FraB (p15A ori, $kan^r$) with a different FraB variant encoded on pET-33b (colE1 ori, $kan^r$), we sought to clone the FraB ORF into a plasmid with compatible origins of replication and antibiotic resistance. To this end, we chose pET-15b (p15A ori, $amp^r$). Cloning of pET-15b–FraB proceeded exactly as previously described for pLANT-2b–$His_6$-TRS-FraB except that FraB_NcoI_F and FraB_BlpI_R were used to amplify FraB from pET-33b–$His_6$-TRS-FraB, BlpI (20 U) and NcoI (20 U) were used for digestion of the amplicons and pET-15b.[4] *Escherichia coli* DH5$\alpha$ ultra-competent cells were used for the transformation and colonies grown on/in LB agar/broth supplemented with 100 µg/mL carbenicillin.[5] Isolation of FraB:$His_6$-FraB heterodimers by nickel-affinity chromatography hinged on saturating amounts of FraB being expressed over $His_6$-FraB such that the formation of $His_6$-FraB homodimers would be negligible and FraB homodimers would not bind to the nickel resin. However, $His_6$-FraB expressed more robustly than FraB rendering this strategy inviable.

High expression was observed for multiple proteins when PA14, a calcium-dependent sugar-binding domain, was fused at the N-terminus.[3] Furthermore, PA14-$His_6$-TRS-FraB:PA14-$His_6$-TRS-FraB (~116 kDa) and $His_6$-TRS-FraB:PA14-$His_6$-TRS-FraB (~96 kDa) dimers are more amenable to separation by size-exclusion chromatography compared to FraB:$His_6$-TRS-FraB (~74 kDa) and $His_6$-TRS-FraB:$His_6$-TRS-FraB (~76 kDa) dimers. We therefore opted to clone pET-16b–PA14-$His_6$-TRS-FraB as described[4] previously for pLANT-2b–$His_6$-SUMO-FraB, albeit with the following modifications. The first PCR reaction for amplifying FraB used FraB_RAIR_F, FraB_RAIR_R, and pET-15b (1 pg) with the following conditions: step 1: 98°C, 2 min; step 2: 98°C, 10 s; step 3: 59-54°C, 15 s; step 4: 68°C, 10 s; step 5: 98°C, 10 s; step 6: 54°C, 15 s; step 7: 68°C, 10 s; step 8: 68°C, 2 min; step 9: 4°C, ∞. To decrease the chances of RAIR transformation with the parental template (pET-15b),[6] consecutive PCR was pursued where a 1:1000 dilution of the first PCR reaction was used as template in a second PCR reaction with otherwise identical composition and conditions.[7] A third and fourth PCR reaction were assembled for amplifying pET-16b–PA14 in a manner analogous to the first and second PCR reactions except using pET16b_F, pET16b_R, and pET-16b–PA14-IBP (courtesy of Dr. Peter L. Davies, Queen's University).[3] The following conditions were used for the third and fourth PCR reactions: step 1: 98°C, 2 min; step 2: 98°C, 10 s; step 3: 65-60°C, 15 s; step 4: 68°C, 63 s; step 5: 98°C, 10 s; step 6: 60°C, 15 s; step 7: 68°C, 63 s; step 8: 68°C, 2 min; step 9: 4°C, ∞. For each PCR reaction, steps 1, 8, and 9 were for one cycle, steps 2-4 were for five cycles where the annealing temperature was decreased by 1°C with each passing cycle, and steps 5-7 were for 25 cycles. Colony PCR was done using FraB_NcoI_F and T7_terminator_R with the following conditions: step 1: 95°C, 5 min; step 2: 95°C, 15 s; step 3: 55°C, 30 s; step 4: 68°C, 64 s; step 5: 68°C, 5 min; step 6: 4°C, ∞. Steps 1 and 6 were for one cycle and steps 2-5 were for 30 cycles. Transformed cells were cultured on/in LB agar/broth supplemented with 100 µg/mL carbenicillin.

Site-directed mutagenesis of pLANT-2b–His$_6$-TRS-FraB and pET-16b–PA14-His$_6$-TRS-FraB proceeded as follows.[4] To generate phosphorylated primers, 10-µL reactions consisting of 1X T4 DNA Ligase Buffer, 4 U of T4 Polynucleotide Kinase, and 8 µM of either FraB_E214A_F and FraB_E214A_R or FraB_H230A_F and FraB_H230A_R were assembled on ice and then incubated at 37°C for 30 min. A 50-µL PCR reaction composed of 1X PrimeSTAR GXL Buffer $Mg^{2+}$ plus, 200 µM dNTP Mixture, 300 nM phosphorylated forward and reverse primers, 0.5 pg/µL of either pLANT-2b–His$_6$-TRS-FraB or pET-16b–PA14-His$_6$-TRS-FraB, 0.05 U/µL PrimeSTAR GXL DNA Polymerase was then assembled in ice. The following PCR conditions were used: step 1: 98°C, 2 min; step 2: 98°C, 10 s; step 3: X°C, 15 s; step 4: 68°C, Y s; step 5: 98°C, 10 s; step 6: Z°C, 15 s; step 7: 68°C, Y s; step 8: 68°C, 2 min; step 9: 4°C, ∞. The "X" and "Z" represent annealing temperatures of 61-56°C and 56°C for E214A and 59-54°C and 54°C for H230A, respectively, while "Y" represents the extension times of 52 s for pLANT-2b–His$_6$-TRS-FraB and 73 s for pET-16b–PA14-His$_6$-TRS-FraB. For each PCR reaction, steps 1, 8, and 9 were for one cycle, steps 2-4 were for five cycles where the annealing temperature was decreased by 1°C with each passing cycle, and steps 5-7 were for 25 cycles. PCR reactions were subjected to DpnI digestion, ethanol precipitation, ligation, and transformation as previously described.[4] All putative positive clones were submitted for Sanger sequencing at the OSUCCC – James Genomics Shared Resource.

Chemically-competent SixPack (Lipinszki et al., 2018)[8] cells were prepared and transformed (with some modification) as described by Yang et al. (2022)[9]. Briefly, 25 µL (50 ng) of pLANT-2b–His$_6$-TRS-FraB and pET-16b–PA14-TRS-FraB in 1X KCM buffer (100 mM KCl, 34 mM $CaCl_2$, 50 mM $MgCl_2$) were mixed with 25 µL thawed cells and incubated on ice for 30 min. Cells were heat shocked at 42°C in a water bath for 90 s, immediately placed back on ice for 2 min, and diluted with 300 µL of room temperature (20°C) SOC prior to being spread on an SOB agar plate containing 35 µg/mL kanamycin and 100 µg/mL carbenicillin. Following an incubation at 37°C for 12 h, single colonies were inoculated into Dynamite media (Taylor et al., 2017) supplemented with 100 mM glucose (in addition to the glucose present in the Dynamite media), 35 µg/mL kanamycin, and 100 µg/mL carbenicillin and incubated at 37°C and 250 rpm for at least 12 h.[10] In all instances, tube/flask : volume ratios that were no more than 1 : 4. Overnight cultures were then diluted 1:100 in Dynamite media (5- and 500-mL for small- and large-scale cultures, respectively) supplemented with 35 μg/mL kanamycin and 100 μg/mL carbenicillin and incubated at 37°C and 250 rpm until cultures reached an optical density (at 600 nm) of 0.6 to 0.8, as determined by a WPA Biowave cell density meter (VWR). Cultures were incubated in a 20°C water bath for 10 min, induced with 1 mM IPTG, and incubated at 20°C and 250 rpm for at least 12 h. Cells were then harvested by centrifugation (2500 g, 15 min) for subsequent purification.

Cell pellets (~2.4 g from 250 mL) were thawed in ice for 30 min and resuspended in 50 mL lysis buffer [20 mM Tris pH 7.5 (20°C), 100 mM NaCl, 2 mM $CaCl_2$, 1 mM DTT] containing a SIGMAFAST Protease Inhibitor Cocktail Tablet (S8830-20TAB, 1003491512; Sigma-Aldrich). A Vibra-Cell Ultrasonic Processor (GEX130) and a large bit (630-0435 P) were used to lyse cells for a total of 5 min: on, 5 s; off, 5 s; 50% amplitude. Lysates were clarified at 30,000 g and 4°C for 15 min and transferred to a separate tube for subsequent purification steps.

Since the PA14 domain binds to dextran-based resins, we used a Sephadex matrix for capturing FraB heterodimers containing the PA14 domain. One gram of Sephadex G-100 (Pharmacia Fine Chemicals, now Cytiva) and 40 mL of Milli-Q water were added to a 50-mL tube and nutated for at least 30 min to allow for resin expansion. Resin was pelleted by centrifugation in a swing-bucket rotor at 500 g for 10 min and the flow-through was discarded. The resin was then twice equilibrated with 40 mL of lysis buffer by nutating for 30 min, centrifuging as described above, and discarding the supernatant. Clarified lysate was added to the resin (in 25-mL batches), nutated at 4°C for 30 min, centrifuged as above, and the flow-through was saved in a separate 50-mL tube. Resin was then washed twice with 25 mL of lysis buffer as described above and washes were transferred to separate tubes. Using lysis buffer to resuspend, the resin was transferred into a 33 (length) by 1.5 (diameter) cm glass column with two, 25-mm glass microfiber filters (Whatman, catalogue #1820-025) placed above the outlet adaptor. Lysis buffer was allowed to drain from column until it was just above the resin, the column was filled with 25% (v/v) elution buffer (lysis buffer + 25 mM glucose), and the inlet adaptor was attached. The column was then attached to an AKTA Purifier FPLC (GE Healthcare, now Cytiva) programmed with a max pressure limit of 0.5 MPa and a max flowrate of 1 mL/min. A gradient of 25 to 100% (v/v) elution buffer, using a total of 4 column volumes (100 mL total), was applied to the column and collected as 1.5-mL fractions.

Aliquots withdrawn from the eluted fractions were subject to SDS-PAGE [15% (w/v) polyacrylamide] at 200 V for 45 min and subsequently visualized with a mixture of glacial acetic acid, methanol, and water (1 : 3 : 6) with (for staining) or without (for destaining) 3 mm Coomassie Brilliant Blue R-250. Fractions containing pure protein were pooled and thrice buffer exchanged (~729-fold total dilution) into lysis buffer using a 10,000 MWCO Amicon® Ultra Centrifugal Filter (Millipore Sigma). Protein concentrations were determined using theoretical extinction coefficients determined by ExPASy ProtParam (SIB Swiss Institute of Bioinformatics). Samples were stored as aliquots at -80°C.

**FraB activity assays**

Activity assays were performed as previously described using fluorescence and an INFINITE M1000 PRO (Tecan, Mannedorf, Switzerland).[11]

**Mass spectrometry**

$His_6$-FraB homodimer and $His_6$-FraB heterodimer variants samples were buffer exchanged into 200 mM ammonium acetate (AmAc) (Sigma-Aldrich) using micro-Bio-spin P6 spin columns (BioRad). The pH was adjusted to 8 using ammonium hydroxide solution (Sigma-Aldrich) to achieve optimal FraB activity.[1] Native MS and SID experiments were performed on Thermo Q Exactive ultra high mass range (UHMR) Orbitrap instrument, which was modified with a surface-induced dissociation (SID) device between quadrupole and C-Trap.[12]

Samples were introduced into the MS instrument by nano-electrospray ionization, using glass capillary tips pulled in-house using a P-97 micropipette puller (Sutter Instruments). 3 to 5 µL of each sample was loaded into the tips, and the electrospray voltage was set between 0.5 – 1.0 kV. The mass spectrometer was operated in positive mode using the following tuning settings: Source temperature: 200°C; detector optimization: low m/z; ion transfer target: high m/z; in-source trapping (IST): 60 V; HCD trap gas flow: 5–7; Resolution: 12,500. SID experiments were performed after isolating the FraB homodimer or heterodimer using quadrupole, with a voltage of 65 V to dissociate the dimer to monomers. Mass spectra were examined using Xcalibur 4.1 (Thermo Scientific), and deconvoluted using UniDec software[13].

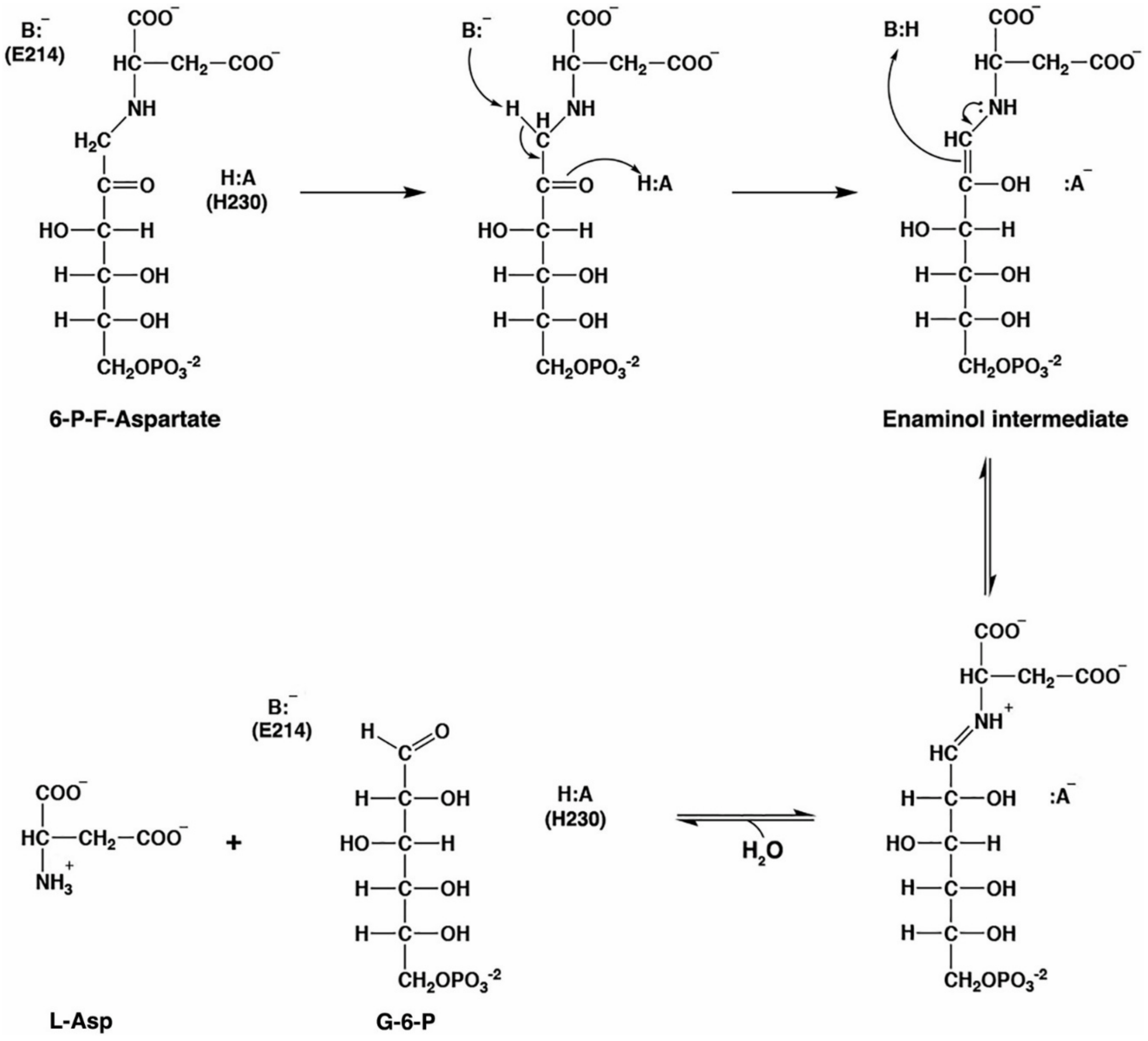


**Figure S1:** A hypothetical catalytic mechanism for FraB. E214 (general base) abstracts one hydrogen from C1 carbon of 6-P-F-Aspartate (6-P-F-Asp, substrate) while H230 (general acid) donates one hydrogen to the carbonyl group to generate an enaminol intermediate. Donation of a proton from E214 to the enaminol intermediate generates a Schiff base that is subsequently hydrolyzed into glucose-6-phosphate (G-6-P) and L-Aspartate (Asp). Reproduced from Sengupta et al. (2019) with no modification.[1]

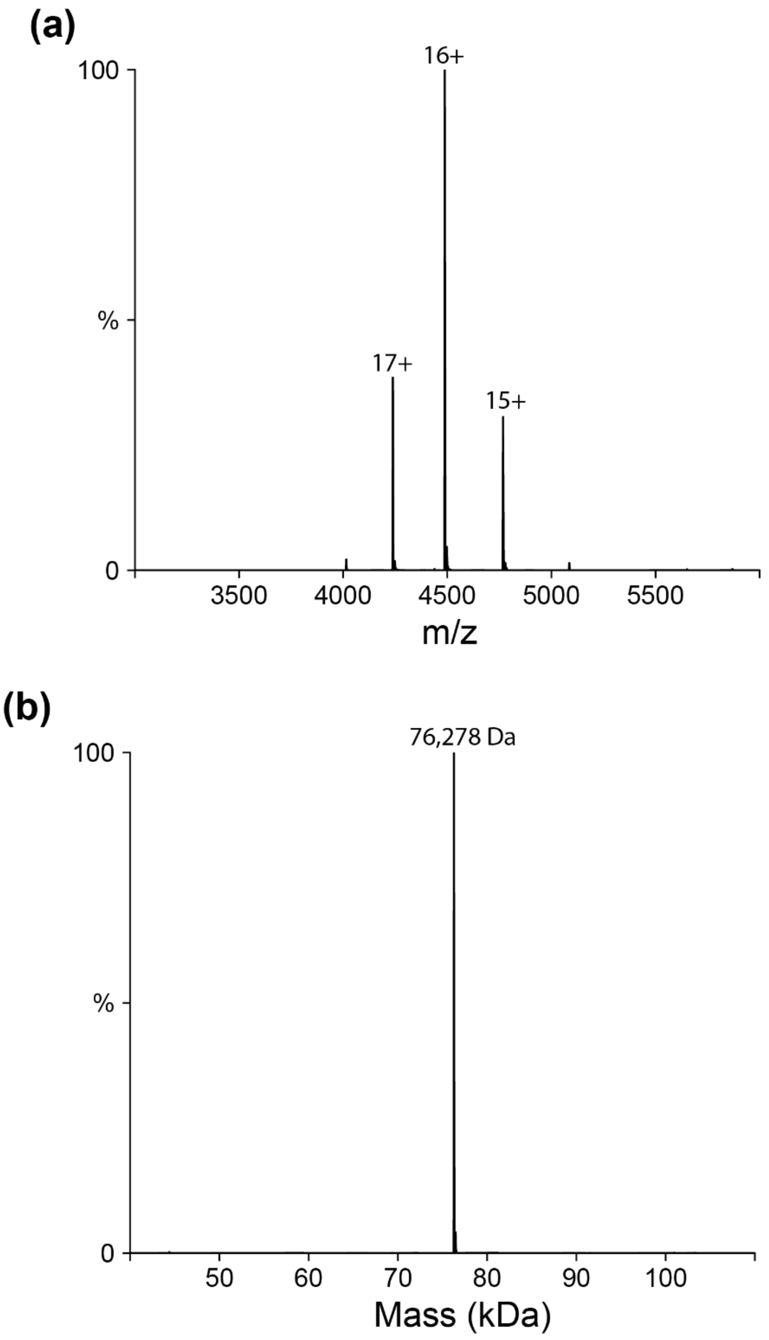


**Figure S2**. Primary (a) and deconvolved (b) mass spectra for $His_6$-FraB homodimer obtained under native conditions (expected, 76,278 Da; observed, 76,278 Da).

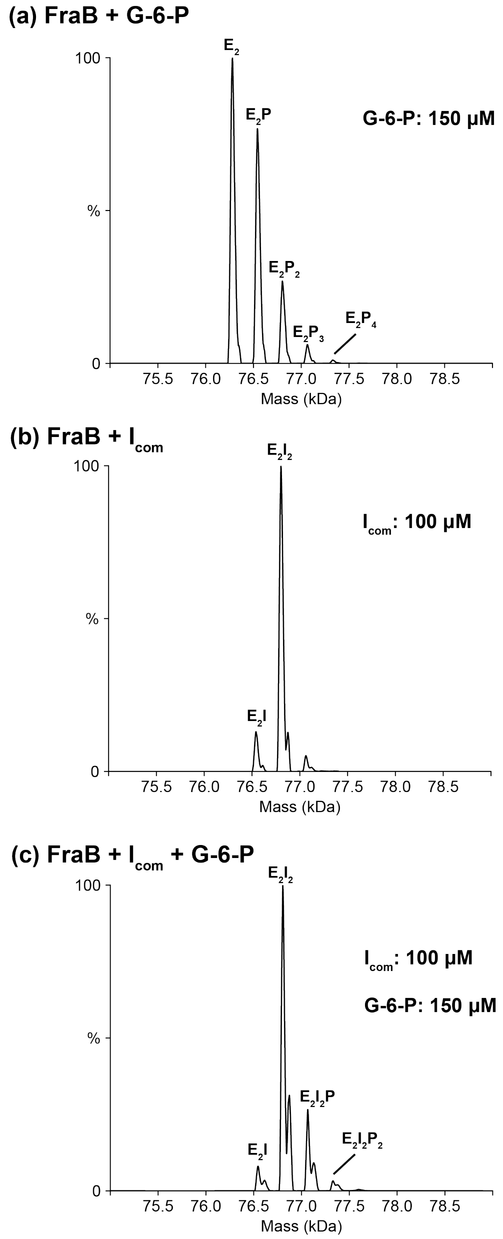


**Figure S3**. Deconvolved native mass spectra for $His_6$-FraB homodimer with product G-6-P (a), competitive inhibitor 77032 (b),[2] and both (c). The final concentrations of FraB, 77032, and G-6-P are 2, 100, and 150 μM, respectively. We tested FraB with free G-6-P (product) and we observed more than two G-6-P copies being bound, a finding that indicates non-specific binding of G-6-P to FraB given a maximum of only two active sites per FraB homodimer. Furthermore, blocking the catalytic sites with the competitive inhibitor 77032 prior to addition of G-6-P generated $E_2I_2P$ and $E_2I_2P_2$, suggesting non-specific binding of G-6-P outside the catalytic pockets. Therefore, the minor $E_2P_2$ population is most consistent with non-specific product rebinding rather than simultaneous catalysis at both active sites.

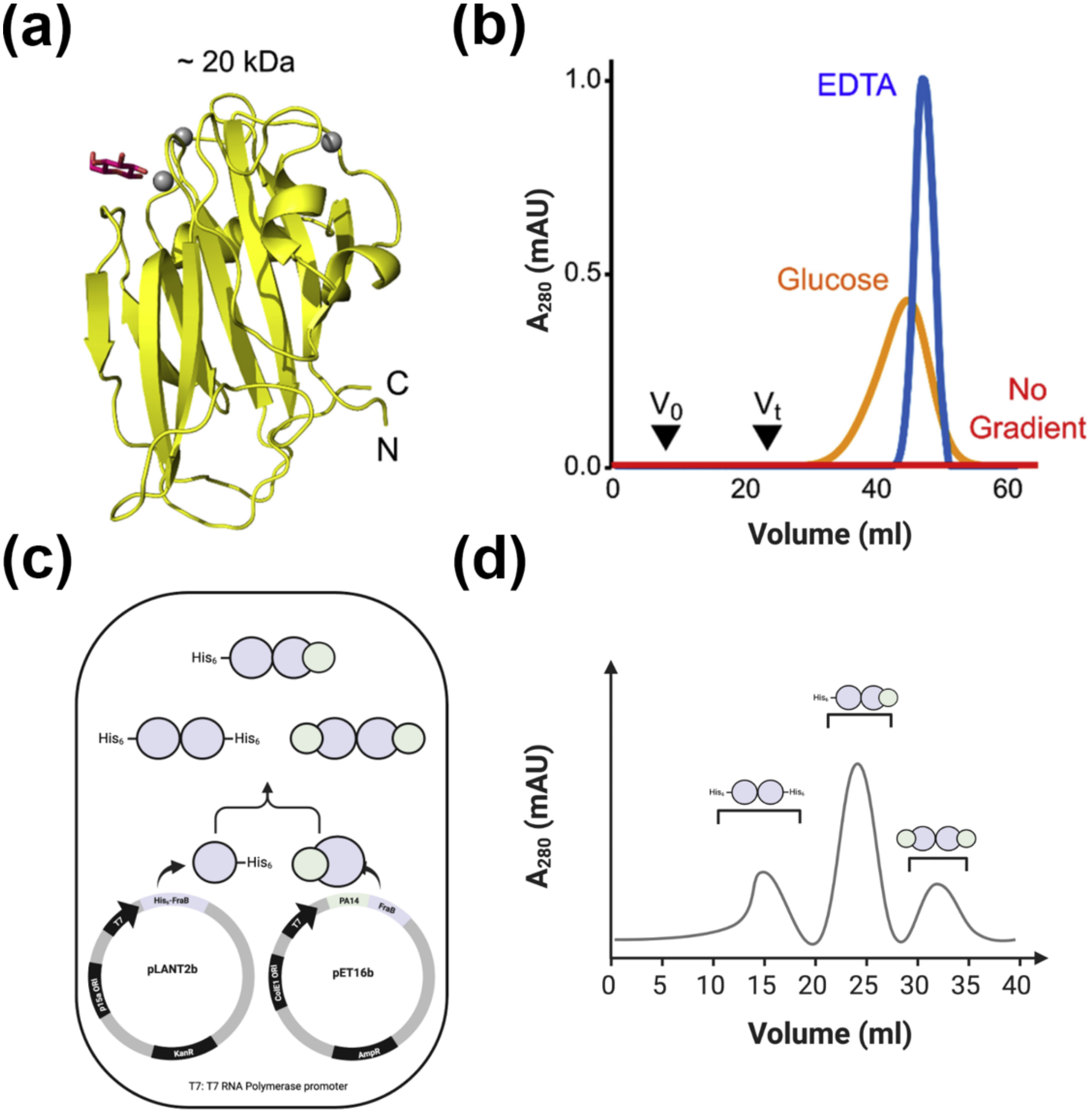


**Figure S4.** The PA14 domain enables economical affinity purification.[3] (a) High-resolution X-ray crystal structure of a bacterial PA14 domain (PDB: 6M8M). Calcium ions (gray) and glucose (red) are depicted. (b) When the PA14 domain is bound to any dextrose resin, it can be eluted with either glucose or EDTA. $V_t$, total column volume; $V_o$, void volume. (c) Schematic showing the expected outcomes of co-overexpression and (d) purification of FraB heterodimers. Panel (a) reproduced with no changes from Kinrade et al. (2020) and panel (b) with minor changes,[3] with permission from Elsevier under license number 6301050944157.

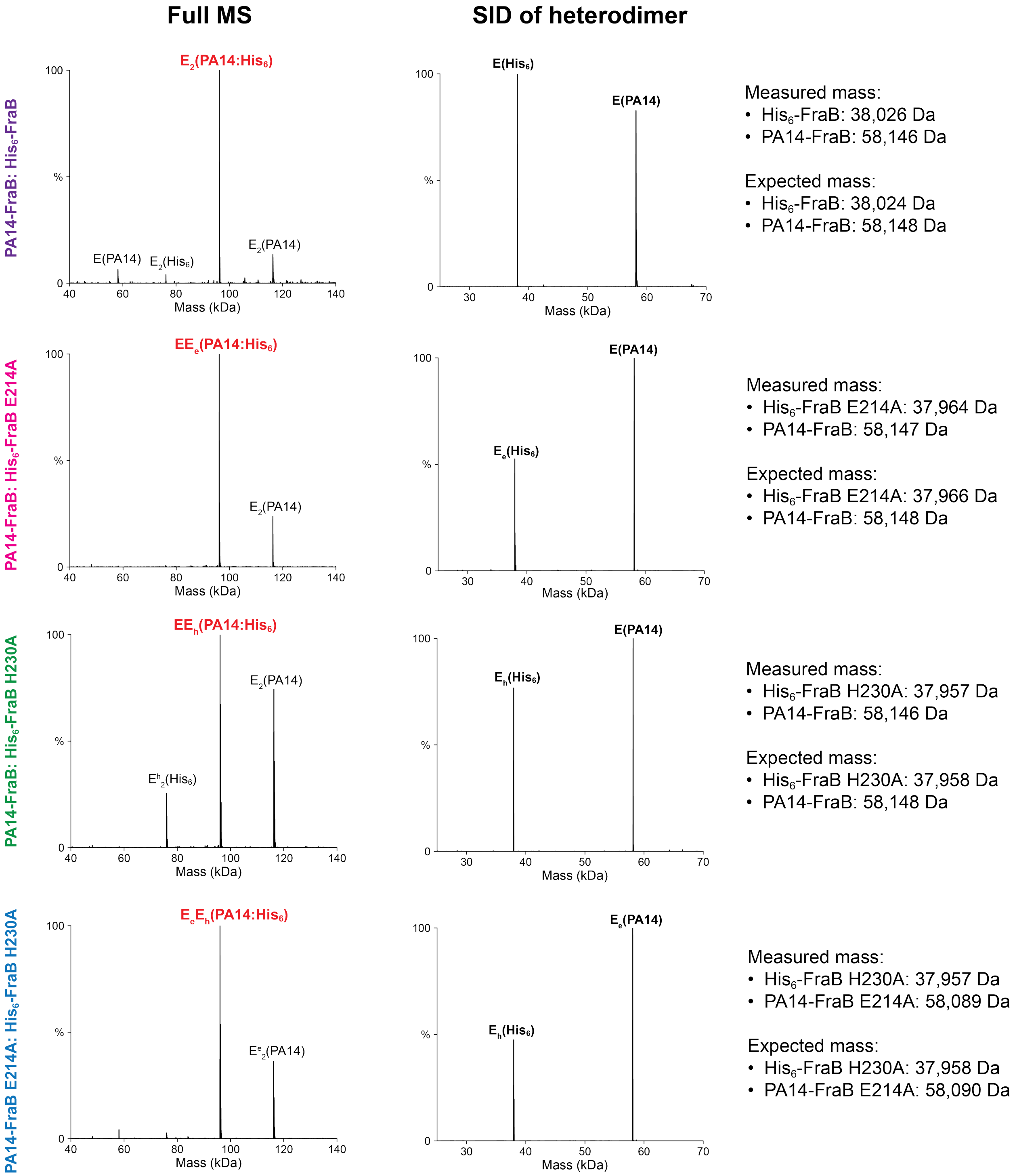


**Figure S5**. Deconvolved full native mass spectra (left) and SID after isolating heterodimer (right), confirming the accurate mass for monomers. All measured masses match the expected masses (< 2 Da error). All SID experiments were at SID 65 V.

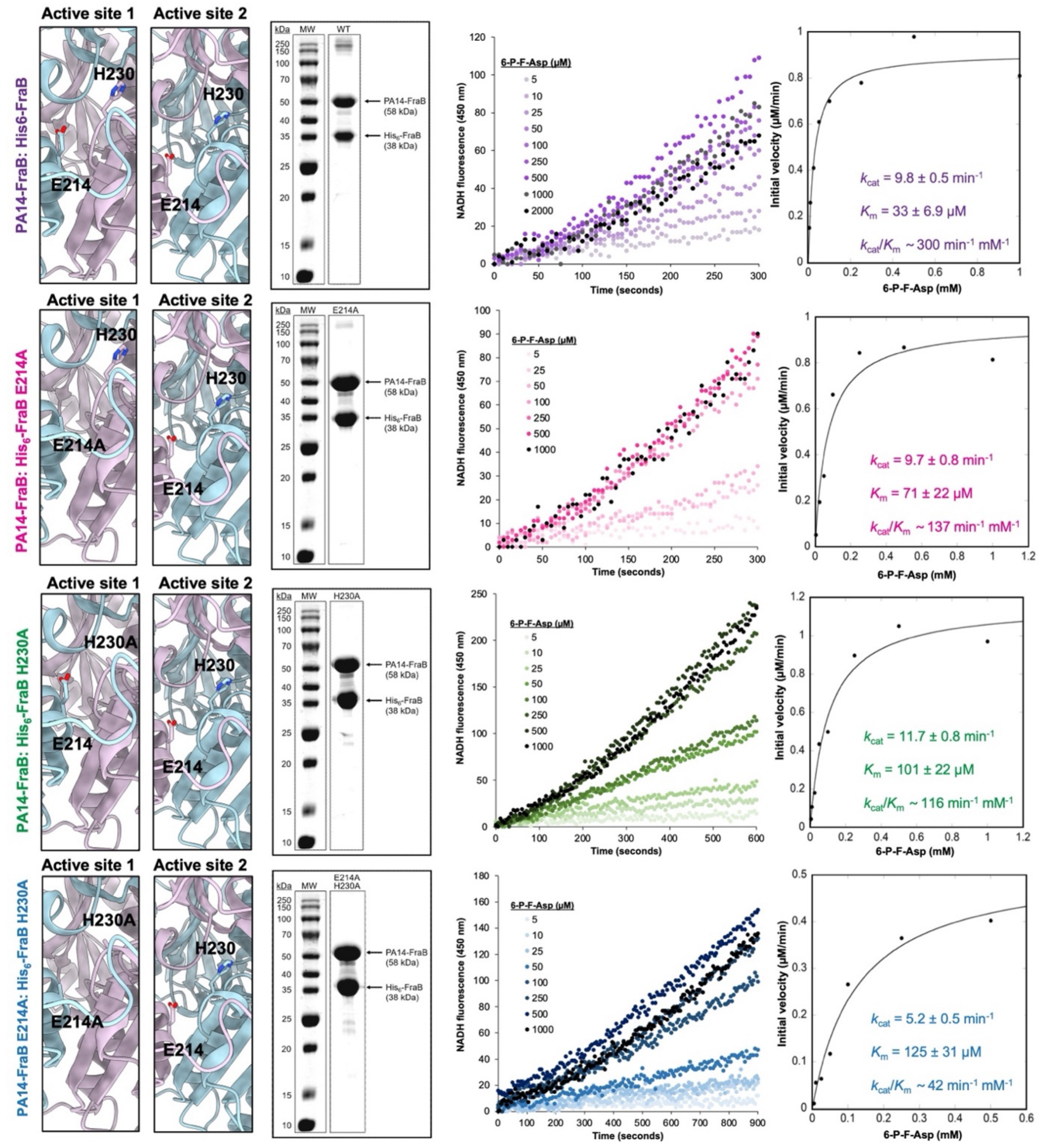


**Figure S6**. Kinetic studies with FraB heterodimer active-site mutants. (Left) Design of the different FraB heterodimer mutants. (Middle-left) Coomassie-stained SDS-15% (w/v) polyacrylamide gels showing purity of the isolated FraB mutants. (Middle-right and right) Primary activity data and Michaelis-Menten analyses for each FraB heterodimer. The correlation coefficient for the Michaelis-Menten curve-fits ranged from 0.97 to 0.99.

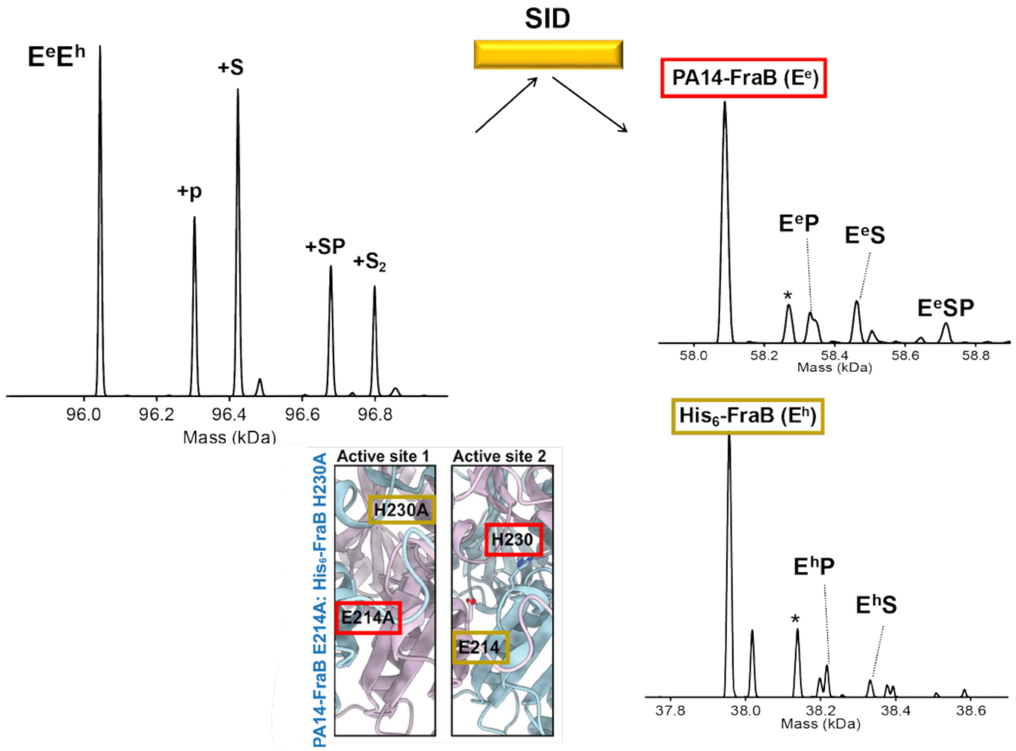


**Figure S7**. SID for FraB heterodimer PA14-FraB E214A:His$_6$-FraB H230A ($E_eE_h$). The heterodimer dissociates into PA14-FraB and His$_6$-FraB monomers while partially retaining substrate and/or product binding. Product- and substrate-bound monomer species (EP and ES) are observed for both subunits, while PA14-FraB additionally exhibits a minor ESP species. Ligand retention on both dissociated subunits is consistent with localization of the active site at the dimer interface. * signifies species modified with AEBSF (+183 Da).[4]

tcgatcccgcgaaattaatacgactcactataggggaattgtgagcggataacaattcccctctagaaataattttgtttaactttaagaa
ggagatatacatatgcatcatcatcatcatcatgaaaacctgtacttccag↓atgatgggtatgaaagagacagttagcaatattgtgac
cagccaggcagagaaaggaggcgttaaacacgtctattacgtggcgtgcggcggttcttatgcggcgttctatccggcgaaagcatttt
tagaaaaagaagcgaaagcgttgactgtcggtctgtataacagcggagaatttattaacaacccgccggtagcgctgggagaaaat
gccgttgtggttgtcgcctcccacaaaggtaatacgccagagacaattaaagcggctgaaatcgcccgtcagcacggcgcgccggt
cattggtttaacctggataatggattcaccgttggtggcgcattgcgactatgtggaaacgtacacgtttggcgacggtaaagatattgcc
ggagagaaaacgatgaaaggcctgctgagtgcggtcgaactgctccagcagacggaagggtatgcgcactacgacgattttcagg
atggcgtcagcaaaatcaaccgtatcgtctggcgcgcttgcgagcaggtagcggagcgtgcgcaggcgttcgcgcaggaatataaa
gacgataaagtcatttataccgtcgccagcggcgcgggctatggcgcagcctacctacagagcatctgcatctttatggaaatgcaat
ggatacattccgcctgtattcatagcggtgagtttttccacgggccgtttgaaattaccgatgcgaatacgcctttcttcttccagttttccga
gggcaatacgcgggcggtggatgaacgcgcgttaaacttcctgaaaaaatatggccgccggattgaagttgtcgatgcgaaagaac
tggggctatcgaccattaaaaccacggttattgattactttaaccactctctctttaataacgtttatcccgtttacaatcgggcgttagctga
ggcgcgtcagcatccgttaacgacgcgccgctatatgtggaaagtggaatattaagctgagcaataactagcataaccccttggggc
ctctaaacgggtcttgaggggttttttgctgaaaggaggaactatatccggatatcccgcaagaggcccggcagtaccggcataacca
agcctatgcctacagcatccagggtgacggtgccgaggatgacgatgagcgcattgttagatttcatacacggtgcctgactgcgttag
caatttaactgtgataaactaccgcattaaagcttatcgatgataagctgtcaaacatgagaattcttagaaaaactcatcgagcatcaa
atgaaactgcaatttattcatatcaggattatcaataccatatttttgaaaaagccgtttctgtaatgaaggagaaaactcaccgaggcag
ttccataggatggcaagatcctggtatcggtctgcgattccgactcgtccaacatcaatacaacctattaatttcccctcgtcaaaaataa
ggttatcaagtgagaaatcaccatgagtgacgactgaatccggtgagaatggcaaaagcttatgcatttctttccagacttgttcaacag
gccagccattacgctcgtcatcaaaatcactcgcatcaaccaaaccgttattcattcgtgattgcgcctgagcgagacgaaatacgcg
atcgctgttaaaaggacaattacaaacaggaatcgaatgcaaccggcgcaggaacactgccagcgcatcaacaatattttcacctg
aatcaggatattcttctaatacctggaatgctgttttcccggggatcgcagtggtgagtaaccatgcatcatcaggagtacggataaaat
gcttgatggtcggaagaggcataaattccgtcagccagtttagtctgaccatctcatctgtaacatcattggcaacgctacctttgccatgt
ttcagaaacaactctggcgcatcgggcttcccatacaatcgatagattgtcgcacctgattgcccgacattatcgcgagcccatttatac
ccatataaatcagcatccatgttggaatttaatcgcggcctcgagcaagacgtttcccgttgaatatggctcataacaccccttgtattact
gtttatgtaagcagacagttttattgttcatgaccaaaatcccttaacgtgagttttcgttccactgagcgtcagaccccgtagaaaagatc
aaaggatcttcttgagatcctttttttctgcgcgtaatctgctgcttgcaaacaaaaaaaccaccgctaccagcggtggtttgtttgccggat
caagagctaccaactctttttccgaaggtaactggcttcagcagagcgcagataccaaatactgtccttctagtgtagccgtagttaggc
caccacttcaagaactctgtagcaccgcctacatacctcgctctgctaatcctgttaccagcgatgataagctgtcaaacatgagaatta
caacttatatcgtatggggctgacttcaggtgctacatttgaagagataaattgcactgaaatctagaaatattttatctgattaataagatg
atcttcttgagatcgttttggtctgcgcgtaatctcttgctctgaaaacgaaaaaaccgccttgcagggcggtttttcgaaggttctctgagct
accaactctttgaaccgaggtaactggcttggaggagcgcagtcaccaaaacttgtcctttcagtttagccttaaccggcgcatgacttc
aagactaactcctctaaatcaattaccagtggctgctgccagtggtgcttttgcatgtctttccgggttggactcaagacgatagttaccgg
ataaggcgcagcggtcggactgaacggggggttcgtgcatacagtccagcttggagcgaactgcctacccggaactgagtgtcagg
cgtggaatgagacaaacgcggccataacagcggaatgacaccggtaaaccgaaaggcaggaacaggagagcgcacgaggg
agccgccagggggaaacgcctggtatctttatagtcctgtcgggtttcgccaccactgatttgagcgtcagatttcgtgatgcttgtcagg
ggggcggagcctatggaaaaacggctttgccgcggccctctcacttccctgttaagtatcttcctggcatcttccaggaaatctccgccc
cgttcgtaagccatttccgctcgccgcagtcgaacgaccgagcgtagcgagtcagtgagcgaggaagcggaatatatcctgtatcac
atattctgctgacgcaccggtgcagcctttttttctcctgccacatgaagcgattcacagatgtctgcctgttcatccgcgtccagctcgttga
gtttctccagaagcgttaatgtctggcttctgataaagcgggccatgttaagggcggttttttcctgtttggtcactgatgcctccgtgtaagg
gggatttctgttcatgggggtaatgataccgatgaaacgagagaggatgctcacgatacgggttactgatgatgaacatgcccggttac
tggaacgttgtgagggtaaacaactggcggtatggatgcggcgggaccagagaaaaatcactcagggtcaatgccagcgcttcgtt
aatacagatgtaggtgttccacagggtagccagcagcatcctgcgatgcagatccggaacataatggtgcagggcgctgacttccgc
gtttccagactttacgaaacacggaaaccgaagaccattcatgttgttgctcaggtcgcagacgttttgcagcagcagtcgcttcacgtt
cgctcgcgtatcggtgattcattctgctaaccagtaaggcaaccccgccagcctagccgggtcctcaacgacaggagcacgatcatg
cgcacccgtggccaggacccaacgctgcccgagatgcgccgcgtgcggctgctggagatggcggacgcgatggatatgttctgcc
aagggttggtttgcgcattcacagttctccgcaagaattgattggctccaattcttggagtggtgaatccgttagcgaggtgccgccggctt

ccattcaggtcgaggtggcccggctccatgcaccgcgacgcaacgcggggaggcagacaaggtatagggcggcgcctacaatcc
atgccaacccgttccatgtgctcgccgaggcggcataaatcgccgtgacgatcagcggtccagtgatcgaagttaggctggtaagag
ccgcgagcgatccttgaagctgtccctgatggtcgtcatctacctgcctggacagcatggcctgcaacgcgggcatcccgatgccgcc
ggaagcgagaagaatcataatggggaaggccatccagcctcgcgtcgcgaacgccagcaagacgtagcccagcgcgtcggccg
ccatgccggcgataatggcctgcttctcgccgaaacgtttggtggcgggaccagtgacgaaggcttgagcgagggcgtgcaagattc
cgaataccgcaagcgacaggccgatcatcgtcgcgctccagcgaaagcggtcctcgccgaaaatgacccagagcgctgccggca
cctgtcctacgagttgcatgataaagaagacagtcataagtgcggcgacgatagtcatgccccgcgcccaccggaaggagctgact
gggttgaaggctctcaagggcatcggtcgacgctctcccttatgcgactcctgcattaggaagcagcccagtagtaggttgaggccgtt
gagcaccgccgccgcaaggaatggtgcatgcaaggagatggcgcccaacagtcccccggccacggggcctgccaccataccca
cgccgaaacaagcgctcatgagcccgaagtggcgagcccgatcttccccatcggtgatgtcggcgatataggcgccagcaaccgc
acctgtggcgccggtgatgccggccacgatgcgtccggcgtagaggatcgagatc

**Figure S8.** The plasmid sequence of pLANT-2b–$His_6$-TRS-FraB with the TEV protease cleavage site indicated by a downward pointing arrow.

gagcaccgccgccgcaaggaatggtgcatgcaaggagatggcgcccaacagtcccccggccacggggcctgccaccataccca
cgccgaaacaagcgctcatgagcccgaagtggcgagcccgatcttccccatcggtgatgtcggcgatataggcgccagcaaccgc
acctgtggcgccggtgatgccggccacgatgcgtccggcgtagaggatcgagatctcgatcccgcgaaattaatacgactcactata
ggggaattgtgagcggataacaattcccctctagaaataattttgtttaactttaagaaggagatataccatggatatggcaccggatgc
acaggcagatagctttggtggtgttgcagttcagggtctgtttggtgaatattatgcctatgcacagggttcagatggtggtaatctgagca
atgttgcacaggttaaagcatttattgcagccaatgaagcagatgccacctttattggtcgcaatattgattatggtagcgttagcggtgat
ctgggtggtaatggtaaagtgcagagctttctgaaagatgatgcaggtagcctgagcaccgatccggaaaattcaagtgatgcaattg
ttaaactgaccggcaatctggaactgcaggcaggcacctatcagtttcgtgttcgtgcagatgatggttatcgtattgaagttaatggtca
gaccgtggccgaatataatggtaatcagggtgcaaatacccgtaccggtagcgaatttaccctgaccggtgatggtccgcatagcgtt
gaaattgtttattggatcagggtggtgcagcacagctgcgtattgaactgcgtgaacaaggtggtgcctatgaaatttttggtagccag
catgcaagccatgggcaccaccaccaccaccacccggaaaacctgtattttcag↓atgatgggtatgaaagagacagttagcaata
ttgtgaccagccaggcagagaaaggaggcgttaaacacgtctattacgtggcgtgcggcggttcttatgcggcgttctatccggcgaa
agcatttttagaaaaagaagcgaaagcgttgactgtcggtctgtataacagcggagaatttattaacaacccgccggtagcgctggga
gaaaatgccgttgtggttgtcgcctcccacaaaggtaatacgccagagacaattaaagcggctgaaatcgcccgtcagcacggcgc
gccggtcattggtttaacctggataatggattcaccgttggtggcgcattgcgactatgtggaaacgtacacgtttggcgacggtaaaga
tattgccggagagaaaacgatgaaaggcctgctgagtgcggtcgaactgctccagcagacggaagggtatgcgcactacgacgat
tttcaggatggcgtcagcaaaatcaaccgtatcgtctggcgcgcttgcgagcaggtagcggagcgtgcgcaggcgttcgcgcagga
atataaagacgataaagtcatttataccgtcgccagcggcgcgggctatggcgcagcctacctacagagcatctgcatctttatggaa
atgcaatggatacattccgcctgtattcatagcggtgagtttttccacgggccgtttgaaattaccgatgcgaatacgcctttcttcttccagt
tttccgagggcaatacgcgggcggtggatgaacgcgcgttaaacttcctgaaaaaatatggccgccggattgaagttgtcgatgcga
aagaactggggctatcgaccattaaaaccacggttattgattactttaaccactctctctttaataacgtttatcccgtttacaatcgggcgtt
agctgaggcgcgtcagcatccgttaacgacgcgccgctatatgtggaaagtggaatattaactcgaggatccggctgctaacaaagc
ccgaaaggaagctgagttggctgctgccaccgctgagcaataactagcataaccccttggggcctctaaacgggtcttgaggggttttt
tgctgaaaggaggaactatatccggatatcccgcaagaggcccggcagtaccggcataaccaagcctatgcctacagcatccagg
gtgacggtgccgaggatgacgatgagcgcattgttagatttcatacacggtgcctgactgcgttagcaatttaactgtgataaactaccg
cattaaagcttatcgatgataagctgtcaaacatgagaattcttgaagacgaaagggcctcgtgatacgcctatttttataggttaatgtca
tgataataatggtttcttagacgtcaggtggcacttttcggggaaatgtgcgcggaacccctatttgtttatttttctaaatacattcaaatatgt
atccgctcatgagacaataaccctgataaatgcttcaataatattgaaaaaggaagagtatgagtattcaacatttccgtgtcgcccttat
tcccttttttgcggcattttgccttcctgtttttgctcacccagaaacgctggtgaaagtaaaagatgctgaagatcagttgggtgcacgagt
gggttacatcgaactggatctcaacagcggtaagatccttgagagttttcgccccgaagaacgttttccaatgatgagcacttttaaagtt
ctgctatgtggcgcggtattatcccgtgttgacgccgggcaagagcaactcggtcgccgcatacactattctcagaatgacttggttgag
tactcaccagtcacagaaaagcatcttacggatggcatgacagtaagagaattatgcagtgctgccataaccatgagtgataacact
gcggccaacttacttctgacaacgatcggaggaccgaaggagctaaccgcttttttgcacaacatgggggatcatgtaactcgccttg
atcgttgggaaccggagctgaatgaagccataccaaacgacgagcgtgacaccacgatgcctgcagcaatggcaacaacgttgc
gcaaactattaactggcgaactacttactctagcttcccggcaacaattaatagactggatggaggcggataaagttgcaggaccactt
ctgcgctcggcccttccggctggctggtttattgctgataaatctggagccggtgagcgtgggtctcgcggtatcattgcagcactgggg
ccagatggtaagccctcccgtatcgtagttatctacacgacggggagtcaggcaactatggatgaacgaaatagacagatcgctgag
ataggtgcctcactgattaagcattggtaactgtcagaccaagtttactcatatatactttagattgatttaaaacttcatttttaatttaaaag
gatctaggtgaagatcctttttgataatctcatgaccaaaatcccttaacgtgagttttcgttccactgagcgtcagaccccgtagaaaag
atcaaaggatcttcttgagatcctttttttctgcgcgtaatctgctgcttgcaaacaaaaaaaccaccgctaccagcggtggtttgtttgccg
gatcaagagctaccaactctttttccgaaggtaactggcttcagcagagcgcagataccaaatactgtccttctagtgtagccgtagtta
ggccaccacttcaagaactctgtagcaccgcctacatacctcgctctgctaatcctgttaccagtggctgctgccagtggcgataagtcg
tgtcttaccgggttggactcaagacgatagttaccggataaggcgcagcggtcgggctgaacggggggttcgtgcacacagcccag
cttggagcgaacgacctacaccgaactgagatacctacagcgtgagctatgagaaagcgccacgcttcccgaagggagaaaggc
ggacaggtatccggtaagcggcagggtcggaacaggagagcgcacgagggagcttccagggggaaacgcctggtatctttatagt
cctgtcgggtttcgccacctctgacttgagcgtcgatttttgtgatgctcgtcagggggggcggagcctatggaaaaacgccagcaacgc
ggcctttttacggttcctggccttttgctggccttttgctcacatgttctttcctgcgttatcccctgattctgtggataaccgtattaccgcctttga

gtgagctgataccgctcgccgcagccgaacgaccgagcgcagcgagtcagtgagcgaggaagcggaagagcgcctgatgcggt
attttctccttacgcatctgtgcggtatttcacaccgcatatatggtgcactctcagtacaatctgctctgatgccgcatagttaagccagtat
acactccgctatcgctacgtgactgggtcatggctgcgccccgacacccgccaacacccgctgacgcgccctgacgggcttgtctgct
cccggcatccgcttacagacaagctgtgaccgtctccgggagctgcatgtgtcagaggttttcaccgtcatcaccgaaacgcgcgag
gcagctgcggtaaagctcatcagcgtggtcgtgaagcgattcacagatgtctgcctgttcatccgcgtccagctcgttgagtttctccaga
agcgttaatgtctggcttctgataaagcgggccatgttaagggcggttttttcctgtttggtcactgatgcctccgtgtaagggggatttctgtt
catgggggtaatgataccgatgaaacgagagaggatgctcacgatacgggttactgatgatgaacatgcccggttactggaacgttgt
gagggtaaacaactggcggtatggatgcggcgggaccagagaaaaatcactcagggtcaatgccagcgcttcgttaatacagatgt
aggtgttccacagggtagccagcagcatcctgcgatgcagatccggaacataatggtgcagggcgctgacttccgcgtttccagactt
tacgaaacacggaaaccgaagaccattcatgttgttgctcaggtcgcagacgttttgcagcagcagtcgcttcacgttcgctcgcgtatc
ggtgattcattctgctaaccagtaaggcaaccccgccagcctagccgggtcctcaacgacaggagcacgatcatgcgcacccgtgg
ccaggacccaacgctgcccgagatgcgccgcgtgcggctgctggagatggcggacgcgatggatatgttctgccaagggttggtttg
cgcattcacagttctccgcaagaattgattggctccaattcttggagtggtgaatccgttagcgaggtgccgccggcttccattcaggtcg
aggtggcccggctccatgcaccgcgacgcaacgcggggaggcagacaaggtatagggcggcgcctacaatccatgccaacccg
ttccatgtgctcgccgaggcggcataaatcgccgtgacgatcagcggtccagtgatcgaagttaggctggtaagagccgcgagcgat
ccttgaagctgtccctgatggtcgtcatctacctgcctggacagcatggcctgcaacgcgggcatcccgatgccgccggaagcgaga
agaatcataatggggaaggccatccagcctcgcgtcgcgaacgccagcaagacgtagcccagcgcgtcggccgccatgccggc
gataatggcctgcttctcgccgaaacgtttggtggcgggaccagtgacgaaggcttgagcgagggcgtgcaagattccgaataccgc
aagcgacaggccgatcatcgtcgcgctccagcgaaagcggtcctcgccgaaaatgacccagagcgctgccggcacctgtcctacg
agttgcatgataaagaagacagtcataagtgcggcgacgatagtcatgccccgcgcccaccggaaggagctgactgggttgaagg
ctctcaagggcatcggtcgagatcccggtgcctaatgagtgagctaacttacattaattgcgttgcgctcactgcccgctttccagtcggg
aaacctgtcgtgccagctgcattaatgaatcggccaacgcgcggggagaggcggtttgcgtattgggcgccagggtggtttttcttttca
ccagtgagacgggcaacagctgattgcccttcaccgcctggccctgagagagttgcagcaagcggtccacgctggtttgccccagc
aggcgaaaatcctgtttgatggtggttaacggcgggatataacatgagctgtcttcggtatcgtcgtatcccactaccgagatatccgca
ccaacgcgcagcccggactcggtaatggcgcgcattgcgcccagcgccatctgatcgttggcaaccagcatcgcagtgggaacga
tgccctcattcagcatttgcatggtttgttgaaaaccggacatggcactccagtcgccttcccgttccgctatcggctgaatttgattgcgag
tgagatatttatgccagccagccagacgcagacgcgccgagacagaacttaatgggcccgctaacagcgcgatttgctggtgaccc
aatgcgaccagatgctccacgcccagtcgcgtaccgtcttcatgggagaaaataatactgttgatgggtgtctggtcagagacatcaa
gaaataacgccggaacattagtgcaggcagcttccacagcaatggcatcctggtcatccagcggatagttaatgatcagcccactga
cgcgttgcgcgagaagattgtgcaccgccgctttacaggcttcgacgccgcttcgttctaccatcgacaccaccacgctggcacccag
ttgatcggcgcgagatttaatcgccgcgacaatttgcgacggcgcgtgcagggccagactggaggtggcaacgccaatcagcaac
gactgtttgcccgccagttgttgtgccacgcggttgggaatgtaattcagctccgccatcgccgcttccactttttcccgcgttttcgcagaa
acgtggctggcctggttcaccacgcgggaaacggtctgataagagacaccggcatactctgcgacatcgtataacgttactggtttca
cattcaccaccctgaattgactctcttccgggcgctatcatgccataccgcgaaaggttttgcgccattcgatggtgtccgggatctcgac
gctctcccttatgcgactcctgcattaggaagcagcccagtagtaggttgaggccgtt

**Figure S9.** The plasmid sequence of pET-16b–PA14-$His_6$-TRS-FraB with the TEV protease cleavage site indicated by a downward pointing arrow. The sequence is color coded as indicated in the name of the construct (PA14-$His_6$-TRS-FraB).

### TABLES

#### Supplementary Table S1

#### DNA oligonucleotides used for cloning in this study

| Primer name | Primer sequence (5’ to 3’) | Primer purpose |
|---|---|---|
| FraB_RAIR_F | ACC CGG AAA ACC TGT ATT TTC AG ATG ATG GGT ATG AAA GAG ACA GTT AG | Amplifying FraB from pET-15b for RAIR cloning into pET-16b |
| FraB_RAIR_R | CAG CCG GAT CCT CGA G TTA ATA TTC CAC TTT CCA CAT ATA GCG G | |
| pET16b_F | CTCGAGGATCCGGCTGCTAACAAAG | Amplifying pET-16b—PA14 for RecA-independent recombination |
| pET16b_R | CTGAAAATACAGGTTTTCCGGGTGG TG | |
| FraB_NcoI_F | ATA TAC CAT GGA TGA TGG GTA TGA AAG AGA CAG TTA G | Amplifying FraB from a parental pET-33b construct and introducing NcoI and BlpI restriction sites at the 5' and 3' ends, respectively, of the PCR amplicon |
| FraB_BlpI_R | TTA TTG CTC AGC TTA ATA TTC CAC TTT CCA CAT ATA GCG G | |
| T7_promoter_F | TAATACGACTCACTATAGGGGAATTG TG | Colony PCR and Sanger sequencing |
| T7_terminator_R | GCTAGTTATTGCTCAGCGG | Sanger sequencing |
| FraB_E214A_F | CTTTATGGCCATGCAATGGATACATT CCG | Mutation of glutamate 214 to alanine |
| FraB_E214A_R | ATGCAGATGCTCTGTAGGTAGG | |
| FraB_H230A_F | GTTTTTCGCGGGGCCGTTTGAAATT ACC | Mutation of histidine 230 to alanine |
| FraB_H230A_R | TCACCGCTATGAATACAGGC | |

**Supplementary Table S2**

**Plasmids used in this study**

| Plasmid | Source | Purpose |
|---|---|---|
| pET-15b | Millipore Sigma (Novagen) | Vector for insertion of FraB |
| pLANT-2b–$His_6$-TRS-FraB | See ref. (4)[4] | Overexpression of $His_6$-TRS-FraB |
| pET-33b–$His_6$-TRS-FraB | See ref. (1)[1] | Source of FraB |
| pET-16b–PA14-$His_6$-TRS-IBP | See ref. (3)[3] | Vector for insertion of FraB |
| pET-15b–FraB | This study | Overexpression of FraB |
| pET-16b–PA14-$His_6$-TRS-FraB | This study | Overexpression of PA14-$His_6$-TRS-FraB |
